# Loss of the benthic life stage in Medusozoa and colonization of the open ocean

**DOI:** 10.1101/2023.02.15.528668

**Authors:** Manon Boosten, Camille Sant, Ophélie Da Silva, Samuel Chaffron, Lionel Guidi, Lucas Leclère

## Abstract

In marine environments, life cycle strategies strongly impact species dispersal and their ability to colonize new habitats. Pelagic medusozoans (jellyfish and siphonophores) exhibit various reproductive strategies, variations of meroplanktonic and holoplanktonic life cycles. In the ancestral meroplanktonic life cycle, a benthic polyp stage alternates with a pelagic medusa stage. During the course of evolution, some medusozoans lost their benthic stage, leading to a holoplanktonic life cycle. The ecological consequences of these losses have not been addressed at global scale. Here, integrating metabarcoding and environmental data from *Tara Oceans* into a phylogenetic framework, we show that each convergent transition toward a holoplanktonic life cycle is associated with a more offshore distribution compared to meroplanktonic medusozoans. Our analyses showed that holoplanktonic medusozoans are more globally distributed and relatively more abundant than meroplanktonic medusozoans, although they are less diversified and occupy a more peripheral position in a global plankton community interactome. This suggests that holoplanktonic medusozoans have acquired a greater tolerance to biotic and abiotic conditions. Overall, our results demonstrate the relationship between medusozoan life cycles, distribution, and biotic interactions, suggesting that the loss of the benthic stage promoted colonization of the open ocean.

## Introduction

Life cycles considerably impact the ecology of organisms^1^. In marine environments, organismal life history strongly affects their dispersal and interactions, and consequently their ecology and evolution^2,3^. Many marine organisms have evolved a complex life cycle, in which adult and larval stages occupy distinct habitats, usually with one stage living in the water column as part of the plankton. The amount of time that organisms spend in the planktonic phase likely impacts their distribution and genetic structure^4–6^. A planktonic life stage is thought to facilitate dispersal, but also to increase the risk of predation, environmental stress, or drifting away from favorable environments^7,8^. Among plankton, the medusozoans (hydrozoans, scyphozoans, cubozoans and staurozoans) represent a ubiquitous^9^ and important component of marine ecosystems through their role in biogeochemical cycles and trophic interactions ^10,11^. Despite their ecological relevance, the main drivers impacting their distribution remain elusive, as most studies are limited to specific taxa or restricted geographic regions, and do not account for the wide diversity of life cycles present in this group.

Most medusozoan species have a complex meroplanktonic life cycle, alternating between benthic and planktonic stages. The ancestral medusozoan life cycle includes a planktonic swimming planula larva, which settles and metamorphoses into a benthic polyp stage that reproduces asexually to give rise to large numbers of sexually mature free-swimming medusae (Fig. 1A). Some medusozoan taxa have lost the benthic stage during evolution and acquired a simplified holoplanktonic life cycle, in which neither embryonic nor adult stages settle on a substrate^12,13^ (Fig. 1A, B). These holoplanktonic life cycles are highly diverse, including species missing the polyp stage (*e.g., Pelagia noctiluca*^14^), species where polyp and medusa forms combine into single colonies (*e.g*., Siphonophorae^15^), and others having a floating polyp stage (*e.g*., Porpitidae). A previous comprehensive literature-based study on hydrozoans attempted to assess the impact of the presence/absence of the benthic stage on the abundance and distribution of species^16^. This study suggested that, on a global scale, holoplanktonic hydrozoan species are less diverse but more widely distributed than meroplanktonic ones. It is currently impossible to generalize these conclusions due to the lack of sufficient comparable *in-situ* data-based studies on medusozoan distribution. This is partly related to sampling difficulties due to their fragile and gelatinous nature^17^, as well as the lack of standardized global data sets and approaches. To bridge the gap, we performed systematic analyses using a comprehensive metabarcoding and environmental data sets generated in the framework of the *Tara Oceans* expedition^18^ (2009-2013), which sampled planktonic organisms across all oceanic regions. By analyzing these data within a phylogenetic context, we investigated the impact of medusozoan life cycles on species distribution, abundance, diversity, and trophic interactions.

**Figure 1:**
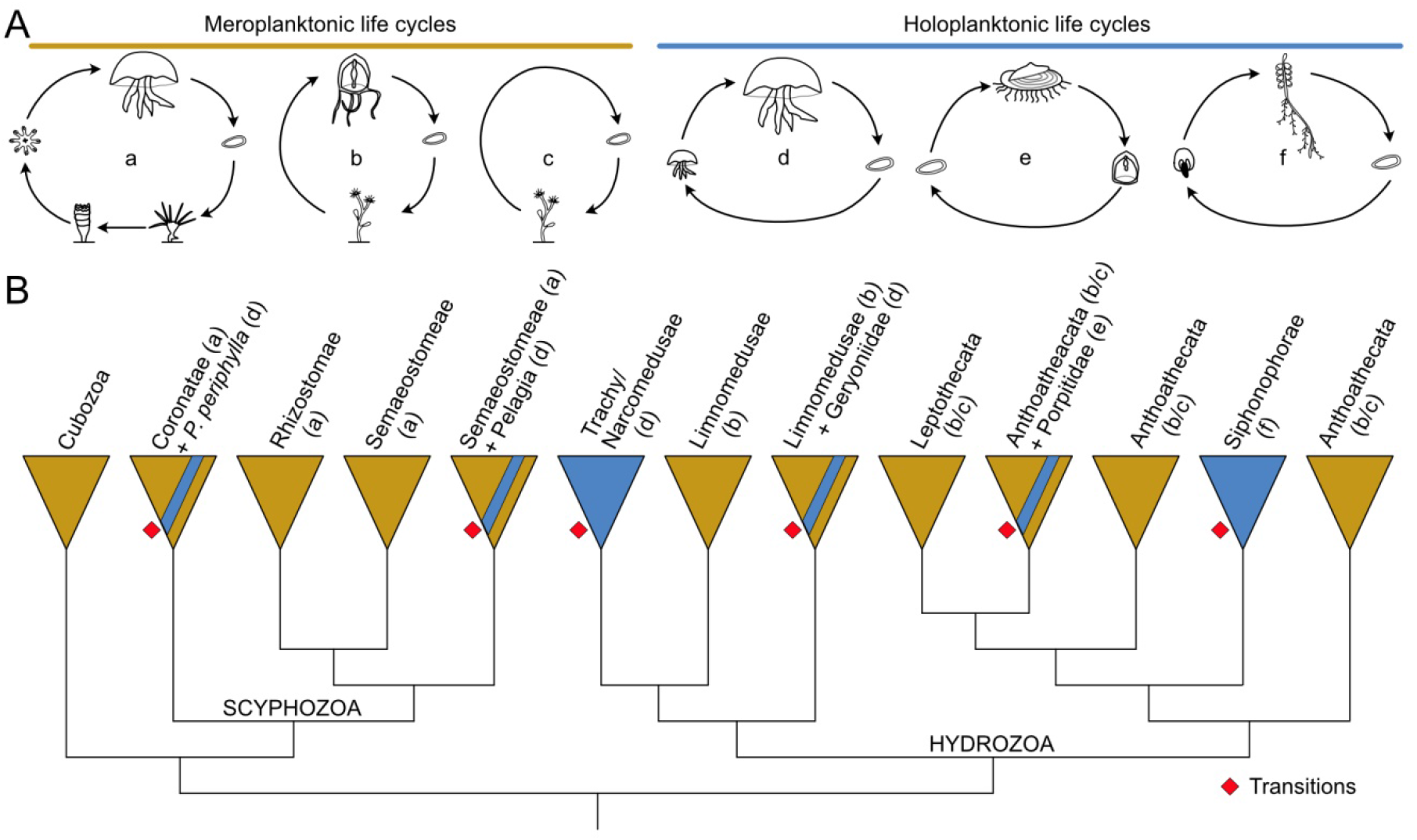
Phylogenetic relationships and life cycle strategies among medusozoans. (A) Schematic representation of the variety of the medusozoan life cycles. Life cycles are represented for a meroplanktonic scyphozoan (a), a hydrozoan with (b) and without (c) medusa stage, holoplanktonic scyphozoan and hydrozoan (d), the holoplanktonic Porpitidae with a planula, a floating polyp and a medusa stage (e) and a siphonophore (f). (B) Simplified phylogeny of medusozoans showing the relationship between the different taxa as well as the different transitions (red squares) from a meroplanktonic to a holoplanktonic life cycle. Some taxa are divided into several clades because they are not monophyletic (e.g Semaeostomae, Limnomedusae and Anthoathecata).

## Results

### Medusozoan life cycle evolution

During the course of evolution, some medusozoan lineages experienced a simplification of their life cycle, including the loss of the benthic stage and the acquisition of a holoplanktonic life cycle^12,14,19^. To determine the pattern of losses, we mapped known life cycle information onto a phylogenetic reconstruction of publicly available medusozoan 18S rRNA genes (Fig. S1A, Table S1). On the basis of the phylogeny and the inferred transitions from a meroplanktonic to a holoplanktonic life cycle, 13 medusozoan taxa were defined (Fig. S1B). We could infer six independent acquisitions of a holoplanktonic life cycle from the ancestral meroplanktonic life cycle, and one possible reversion from a holoplanktonic to a meroplanktonic life cycle in the siphonophore Rhodaliidae. Of these acquisitions, four occurred within Hydrozoa, notably in the common ancestors of Trachymedusae/Narcomedusae, Siphonophorae, Geryoniidae, and Porpitidae. In scyphozoans, at least two lineages lost their benthic stage, *Pelagia* and *Periphylla periphylla* (Fig. 1B). Using calibration points (Table S2), we could date the transitions from a meroplanktonic to a holoplanktonic life cycle. The oldest transition from a mero- to a holoplanktonic life cycle was inferred in the common ancestor of Siphonophorae during Ordovician or Silurian, followed by the Geryoniidae and the Trachymedusae/Narcomedusae. The most recent transition occurred in the *Periphylla* lineage during Paleogene or Neogene (Fig. 2A). By relating these transitions to the relative sea level and continent fragmentation index, we noted that the majority of the transitions occurred at a time when the relative sea level was high and continents fragmented, while no transition occurred between the late Devonian and Triassic where the sea level was lower and the continent merged into the supercontinent Pangea (Fig. 2B).

**Figure 2:**
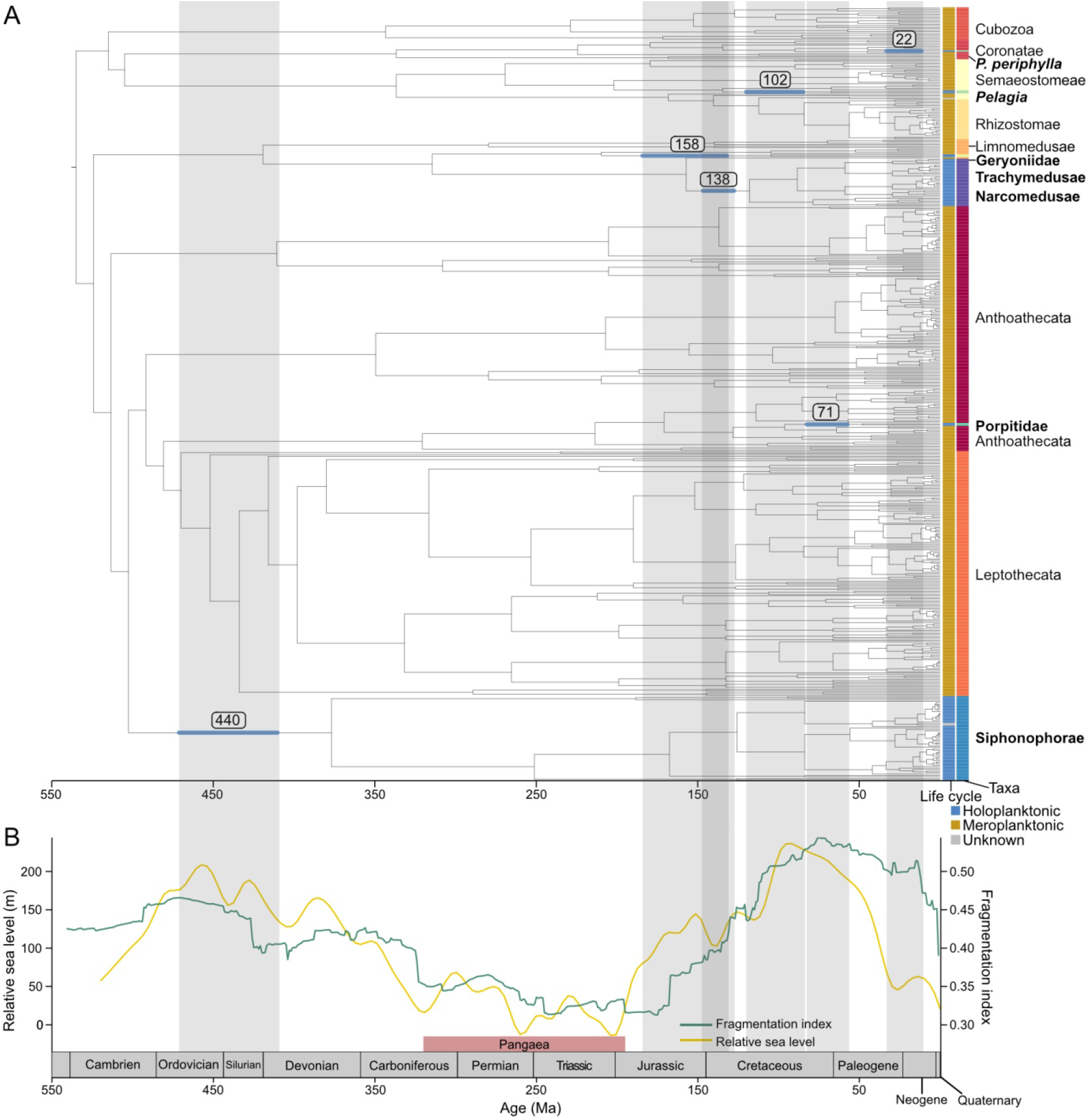
Dating of transitions from a meroplanktonic life cycle to a holoplanktonic life cycle, in relation with the relative sea level. (A) 18S rDNA maximum-likelihood time-calibrated medusozoan species tree with geological time-scale in million years. The tree was rooted with the Anthozoa; holoplanktonic taxa are written in bold. The average (squared value), first and third quartile (blue line) of the age of each of the six transitions from a meroplanktonic to a holoplanktonic life cycle are indicated on the tree. (B) The relative sea level changes from Marcilly et al. (2022)^73^ and the fragmentation index from Zaffos et al. (2017)^42^ through Earth history.

### Medusozoan composition in the *Tara Oceans* data set

In this study, we explored the large-scale and global plankton metabarcoding data sets from the *Tara Oceans* expedition (2009-2013). The sampling strategy of planktonic organisms was based on size fractionation to recover a wider variety of organisms. In the 210 sampling stations, samples were collected with different plankton net mesh sizes and at different depths (surface: 5 meters (m) average depth; deep chlorophyll maximum: 17-188 m), covering a large variety of environments. All samples were systematically processed for amplicon sequencing and analyses, which allowed the generation of high-quality and standardized data sets^18,20,21^. To compare the diversity and abundance of medusozoan taxa across the *Tara Oceans* samples, we first extracted medusozoan Operational Taxonomic Units (OTUs) from the 18S V9 rRNA metabarcoding data set. We mapped them onto our 18S rRNA medusozoan phylogeny to infer the taxa and the life cycle for each medusozoan OTU (*i.e*., meroplanktonic or holoplanktonic) (Fig. S1A). After filtering on the basis of the relative abundance and the number of samples where OTUs were detected (see Methods), 300 medusozoan OTUs were finally retained. These filters allowed to control for false positives and did not impact the sampled taxa diversity (Fig. S1B). We then performed a series of analyses to show that the assembled data set covers a comprehensive fraction of medusozoan life cycles and diversity. All selected OTUs were at least assigned at the order level; 196 could be assigned at the family level, and 32 at the species level. Among the medusozoan OTUs, 182 were inferred as meroplanktonic medusozoan (MP), and 118 as holoplanktonic medusozoan (HP) (Table S3). Of the 13 taxa previously defined, 11 of these taxa were represented by at least one OTU in our filtered *Tara Oceans* data set (Fig. S1B). 92% of the OTUs belonged to Hydrozoan (277 OTUs) and 8% to Scyphozoa (23 OTUs), and none to Cubozoa or *Periphylla* (a *Periphylla periphylla* OTU was excluded, as indistinguishable from the meroplanktonic *Paraphyllina ransoni* OTU). The two most species-rich medusozoan taxa, the Leptothecata and Anthoathecata^22^, were also the best-represented in our data set, totaling 101 OTUs and 62 OTUs respectively (Fig. S1C-F). Among the 814 plankton samples (station/depth/fraction) of the *Tara Oceans* data set, at least one medusozoan OTU was detected in 781 of them. Medusozoan relative abundances were higher in the largest size fraction (180-2000 μm) in comparison with the other size fractions (Fig. S2) with still 13% of the medusozoan reads found in the smallest size fraction sampled (0.8-5μm), which could correspond to gametes or small pieces detached from mature specimens (Table S4). Rarefaction analyses of MP and HP OTUs were performed by measuring both their richness (number of OTUs) and diversity (Shannon index) as a function of the number of samples. Both metrics showed saturation at the global scale and higher values for MP compared to HP OTUs (Fig. S3). Thus, the assembled data set covers a comprehensive fraction of medusozoan life cycles and diversity and allows addressing questions about medusozoan distribution, diversity, and potential biotic interactions.

### Distribution and diversity of medusozoan OTUs

The diversity of MP and HP OTUs was estimated using the Shannon Index by station, by averaging the relative abundance of OTUs in all the samples from the same station (depth/fraction, see Methods). MP and HP OTUs alike showed a latitudinal gradient with maximum diversity in the tropical and subtropical regions and minimum diversity in the pole (Fig. 3A). Diversity values were significantly higher for MP OTUs than for HP OTUs (*Wilcoxon-test p* = 1.4 x 10^-4^, Fig. 3A). The relative abundance showed instead an inverse trend with higher values for HP OTUs (ranging from 1.7 x 10^-5^ % to 44%) than MP ones (ranging from 7 x 10^-6^ % to 26%), notably in the tropical and subtropical regions (Fig. 3B, Fig. S4A). In 91% of samples collected during the *Tara Oceans* expedition, the proportion of reads belonging to HP OTUs was higher than MP OTUs. Some exceptions were found in the Arctic and for locations close to the shore, such as in South Africa (treated extensively below, Fig. 3C, Table S5).

**Figure 3:**
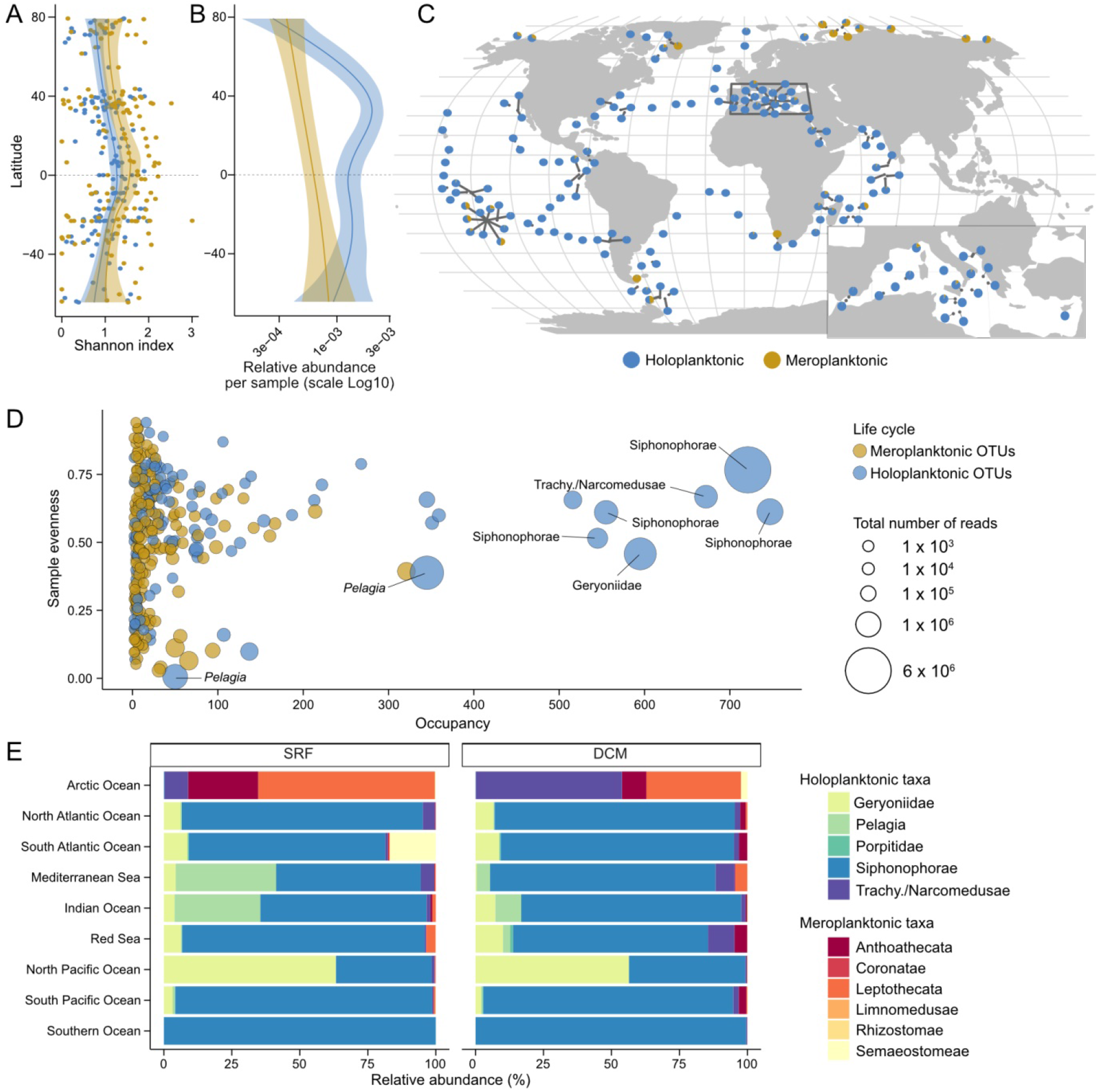
Global distribution and diversity of medusozoans. (A) Latitudinal gradient diversity for both types of life cycle by station determined with the Shannon index; the relative abundance of OTUs in the samples (depth/fraction) from the same station were averaged. (B) Latitudinal gradient of relative abundance for both types of life cycle for the SRF samples 180-2000μm with the x-axis log10 scale. (C) World Map showing the proportion of relative abundance of holoplanktonic (blue) and meroplanktonic (yellow) medusozoan OTUs in each *Tara Oceans* surface sample for the largest size fraction available in the station. (D) Occupancy, evenness and abundance of each medusozoan OTU among samples. A circle represents a medusozoan OTU according to its occupancy (x-axis) and evenness (y-axis) among samples. Occupancy represents the number of samples in which the OTU is detected; evenness represents the homogeneity of the distribution over the samples where the OTU is present. The size of the circle indicates the total number of reads per OTU. (E) The proportion of relative abundance of OTUs assigned to the different medusozoan taxa according to Longhurst biomes Longhurst (2010)^63^ for samples 180-2000μm.

The medusozoan community was globally dominated by a few abundant and widespread HP OTUs (Fig. 3D) belonging to Siphonophorae, Trachymedusae, Narcomedusae, Geryoniidae, and *Pelagia*, with a maximum occupancy of 747 over 781 samples for a calycophoran siphonophore OTU. MP OTUs were instead less prevalent, with a maximum occupancy of 321 for an Anthoathecata OTU (Fig. 3D). The same trend emerged from the medusozoan composition of each oceanic region with the three HP taxa; Siphonophorae, Geryoniidae, and *Pelagia*, representing respectively on average 64%, 9%, and 8% of the relative abundance (Fig. 3E, Fig. S4B). Conversely, the Arctic Ocean was dominated by MP OTUs belonging to the Leptothecata, Anthoathecata, and Semaeostomeae (Fig. 3E). Overall, HP taxa, despite being less diversified than MPs, displayed a higher relative abundance at global scale, thus raising questions about biotic factors influencing their distribution.

### Potential biotic interactions of medusozoans

To assess the place of MPs and HPs in a plankton ecological network, we investigated the co-abundance of medusozoan OTUs within a global plankton association network^23^. This interactome was constructed from *Tara Oceans* metabarcoding data and inferred potential biotic interactions from OTUs co-abundance profiles (Fig. 4A). Of the 300 OTUs selected in the previous analyses, 153 medusozoan nodes were found in the interactome with 100 HP nodes and 53 MP nodes. To investigate the position of medusozoans in the network, two network topological metrics were computed: the node degree^24^ (number of edges connected to a node), and the betweenness centrality^25^ (number of shortest paths passing through the node). The comparison of graph topological metrics (Fig. 4B) showed that the HP nodes had a significantly lower node degree and betweenness than the MP nodes (Wilcoxon-test *p* = 0.0012 and 0.0031), suggesting a more peripheral place of the HPs in the interactome. Further, HP and MP nodes mainly co-occurred with Alveolata (32.3% of the HP nodes edges, 32% of MP nodes edges), Rhizaria (10.5% and 13.8% respectively), non-medusozoan metazoans (8.4%, 9% respectively), and other medusozoans belonging to the same life cycle category (11% for the HP and 7% for the MP). Regarding associations between HP and MP nodes, we detected few edges between HP and MP nodes (2%, Fig. 4C). Additionally, only a few non-medusozoan nodes co-occurred with both HP and MP nodes (24 nodes), and overall HPs and MPs co-occurred with the same broadly defined taxa, even if the specific OTUs were distinct. All five previously defined interactome communities (TC0: Trades-like, PC1: Polar-like, WC2: Westerlies-like, TC3: Trades-like, UC4: Ubiquitous)^23^ included medusozoan nodes, but did not display the same medusozoan community composition (Fisher test: *p*-value = 5 x 10^-4^, Fig. S5). Three out of the five communities were strongly dominated by HPs (WC2: 82.3% HP, TC3: 83.3% HP, UC4: 100% HP), and two were composed of a balanced proportion of HPs and MPs (TC0: 47.7% HP, PC1: 51.3% HP). Thus, HPs appeared overall less central than MPs in the plankton interactome, probably highlighting that HPs colonized distinct niches and biomes while both co-occurring with different sets of OTUs.

**Figure 4:**
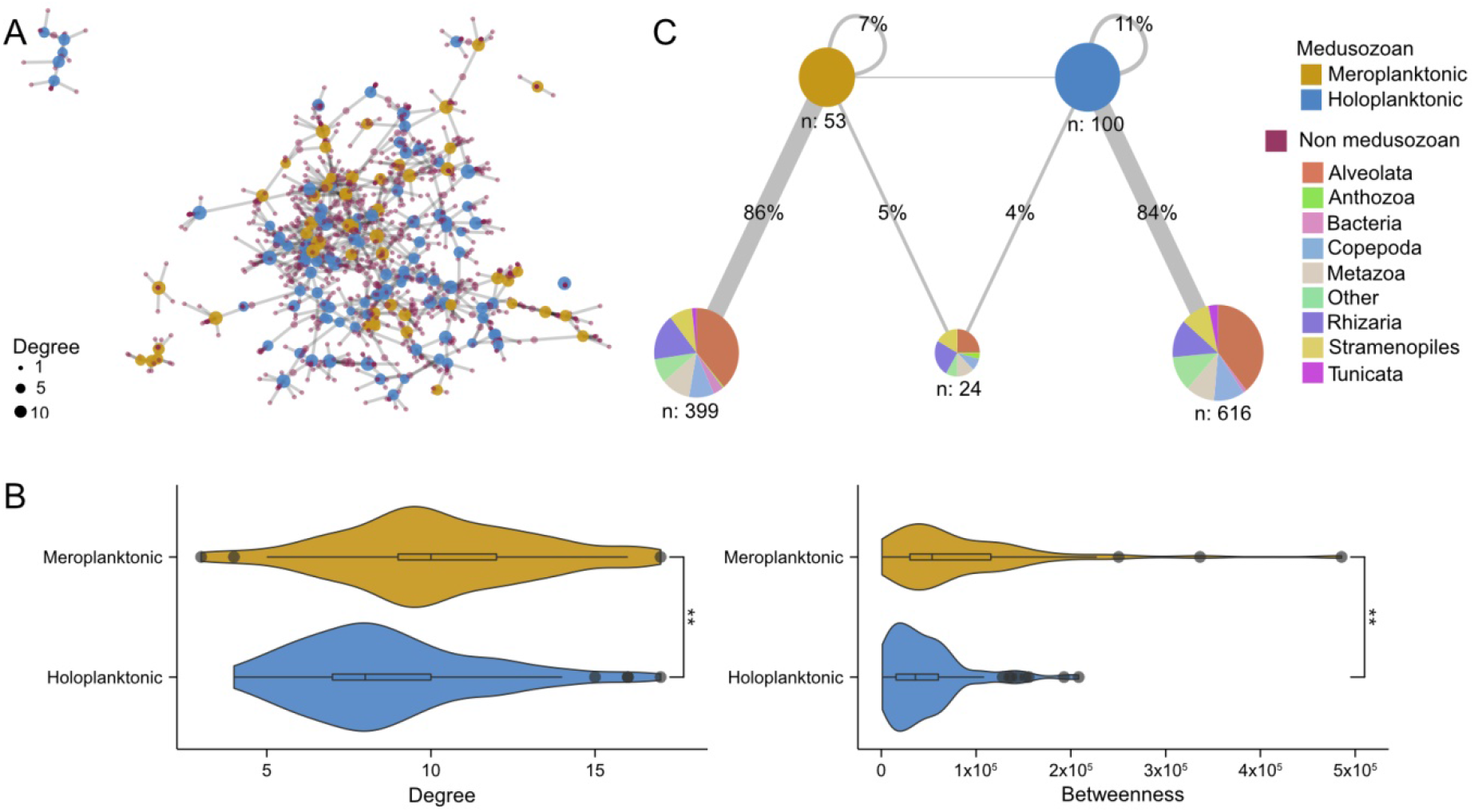
Place of the medusozoans in the plankton interactome^23^. (A) Representation of the medusozoan and their direct neighbors network topology. HP nodes are colored in blue, MP nodes in yellow and non medusozoan nodes in purple, the size of the dots is proportional to their degree. (B) Degree and betweenness, calculated on the entire interactome, of medusozoan nodes according to their type of life cycle. *P*-values were computed using a Wilcoxon test. ** *p*-value <= 0.01. (C) Simplified co-occurrence network of medusozoan nodes and their direct neighbors. Nodes are merged into 5 representative pie charts: HP nodes, MP nodes, direct neighbors of both HP and MP nodes, direct neighbors of only HP nodes, and direct neighbors of only MP nodes. n corresponds to the number of nodes in each represented pie chart. The proportion of positive correlation edges linking HP or MP nodes to each of the five representative pie charts is indicated in percentages.

### Environmental factors impacting medusozoan distribution

A redundancy analysis (RDA) was performed to determine to what extent the detected variation in medusozoan composition among samples could be explained by environmental factors. The RDA on all oceanic regions showed that most of the variance was indeed explained by distinct environmental settings in the Arctic (Fig. S6A), mostly dominated by MP medusozoans (Fig. S6B). Thus, the Arctic samples were excluded from further analyses. Besides the Arctic region, we did not detect clear region-specific medusozoan community, or differences between size fractions (Fig. S6C). Some taxa were not associated to environmental variables in their distribution (Rhizostomae, Porpitidae, Limmnomedusae, Trachymedusae/Narcomedusae) while others were clearly linked (Semaeostomeae, Siphonophorae, Geryoniidae, Leptotecata; Fig. 5A). The two first axes of the RDA explained a small proportion of the variance (~10 % in total). The first RDA axis (5.9% of variance explained) strongly contrasted a major cluster of samples dominated by HP taxa (blue dot, RDA1 > 0) with samples that were dominated by MP taxa (yellow dots, RDA1 < 0). This axis was positively correlated with the bathymetric depth (henceforth BD), and was negatively correlated with turbidity, expressed as the absorption coefficient of the colored dissolved organic matter (acCDOM) and the particle backscattering coefficient of particles at 470 nm (bbp470). The second RDA axis (4%) appears to mark an opposition between samples from dynamic eutrophic zones and samples coming from less dynamic zone (high residence time). This second axis, which was positively correlated to the nitrate and photosynthetically active radiation (PAR) and negatively correlated to the residence time, did not explain any variation in medusozoan composition according to the life cycle. In conclusion, the first RDA axis distinguished the coastal environments, characterized by shallow and turbid water, from the deeper and less turbid open oceans. The HP OTUs were found in samples with a statistically higher BD and lower turbidity in comparison with the MP OTUs (Fig. 5B). This suggests that medusozoan distribution is spatially organized, according to the life cycle. The HPs comprised the majority of the total medusozoan relative abundances across the BD gradient, with their proportion gradually increasing with increasing depth (Fig. 5C, Fig. S7A). The proportion of the samples dominated by the MPs was the highest in the shallowest samples (0-200 m; 35%), then decreased between BD 200-500 m to 27% and dropped to less than 1.3% with BD below 500 m (Fig. 5B). The same trend was observed when considering the distance from the coast with the majority of samples collected at more than 200 kilometers from the coast dominated by HPs (Fig. S7B). Thus, samples dominated by HP OTUs were mainly in the open ocean, while samples dominated by MP were mainly coastal.

**Figure 5:**
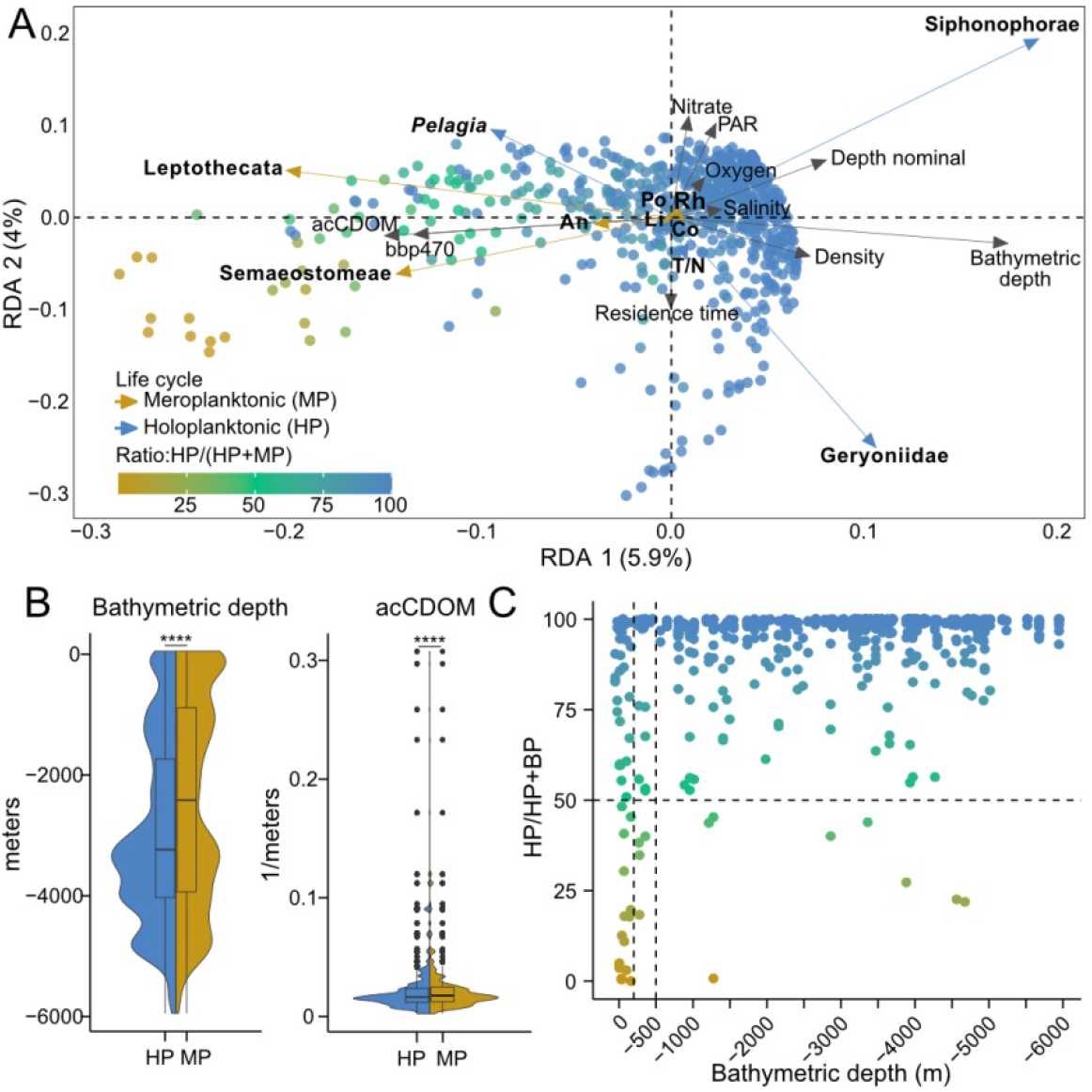
Impact of environmental variables on medusozoans distribution, excluding the Arctic samples. (A) Triplot of the redundancy analysis (RDA) showing the explanatory environmental variables in black, and the response variable, the relative abundance of medusozoan taxa, in blue for HP taxa and yellow for MP taxa. Each dot corresponds to a sample (i.e. one station at one depth and one size fraction) and is colored by the life cycle ratio (abundance of HP OTUs on total medusozoan abundance) so that the blue dots represent samples dominated by HP OTUs and the yellow dots samples dominated by MP OTUs. Abbreviations: An Anthoathecata, Co Coronatae, Li Limnomedusae, Po Porpitidae, Rh Rhizostomae, T/N Trachymedusae/Narcomedusae. The adjusted R-squared of the RDA is 0.13 (0.12 unadjusted), with 9.9. % of the variance explained by the first two axes (B) Violin and boxplot plot reporting the bathymetry and the acCDOM-absorption coefficient of colored dissolved organic matter-values for each OTUs according to the type of life cycle. *P* values were computed using Wilcoxon test, **** *p*-value <= 0.0001. (C) Bathymetric depth according to the life cycle ratio in the samples each dot represents a sample colored by the ratio. The first vertical line corresponds to −200m and the second to - 500m.

### Environmental changes associated with the acquisition of a holoplanktonic life cycle

To further investigate the relationship between life cycles and environmental variables, we used a phylogenetic approach. A medusozoan OTUs phylogenetic tree was constructed by extracting OTUs placement in the phylogenetic reconstruction of publicly available medusozoan 18S rRNA genes (see Methods). Median BD for each OTU was then mapped onto this tree. The reconstruction highlighted that each independent acquisition of a holoplanktonic life cycle corresponds to an increase in the median BD (Fig. 6A). Furthermore, each taxon that independently lost the benthic stage showed statistically higher BD values than its closest related meroplanktonic taxon (Fig. 6B). To statistically test the relationship between the environmental variables and the type of life cycle, a phylogenetic logistic regression was performed (Fig. 6C). It showed a strong relationship between the median BD and the type of life cycle (*p*-value = 2.6 x 10^-15^), suggesting that the life cycle influences the type of environment colonized by medusozoans. Analogous mapping of the median turbidity, instead, did not support a solid correlation between the acquisition of a holoplanktonic life cycle and a decrease in turbidity (Fig. S8A). Phylogenetic logistic regression (Fig. S8B) showed a weaker relationship between the type of life cycle and median turbidity (*p*-value = 2.5 x 10^-5^) in comparison with the BD. This correlation seems to be mainly driven by some Anthoathecata and Leptothecata meroplanktonic clades found in a higher turbidity (acCDOM), not related to the repeated losses of the benthic stage in Medusozoa (Fig. S8A). Taken together, these results suggest that the loss of the benthic stage has likely impacted environments colonized by medusozoans, with the BD as the most discriminating variable between meroplanktonic and holoplanktonic medusozoans.

**Figure 6:**
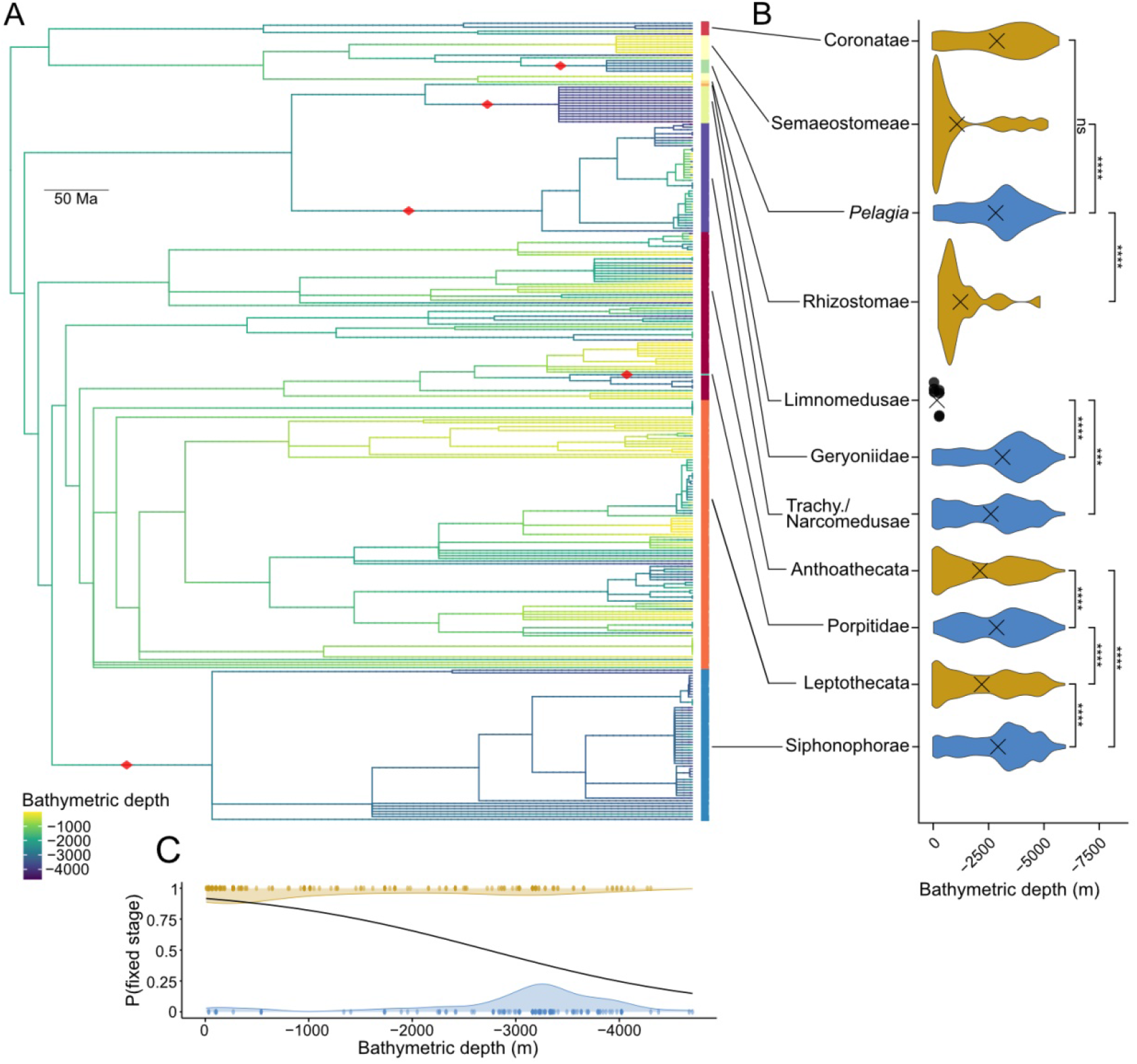
Phylogenetic relationship between medusozoan life cycles and the bathymetric depth. (A) Phylogenetic tree of OTUs on which the evolution of the median value of bathymetric depth has been estimated using maximum likelihood and used to color the branches of the tree. The red square illustrates transitions from a meroplanktonic life cycle to a holoplanktonic life cycle. (B) Violin plot reporting the bathymetric depth values by OTUs per taxa. P-value were computed using a Wilcoxon test, **** p-value <= 0.0001;*** p-value <= 0.001; ** p-value <= 0.01; ns not significant. (C) Phylogenetic logistic regression showing the relationship between the bathymetric depth and the type of life cycle (p-value: 2.6 × 10-15).

## Discussion

The *Tara Oceans* expedition provides standardized data about the planktonic environment on a global scale, which allows to uniquely compare oceanic regions^18,20^. Using this comprehensive data set, we showed that medusozoans (both HPs and MPs) display a latitudinal diversity gradient (Fig. 3C), which is a widely observed pattern across both terrestrial and marine organisms^21,26^. We further detected a coastal to open ocean gradient in medusozoan distribution, with MPs preferentially located in coastal waters, while HPs being present both in the coastal and open ocean waters. Bathymetric depth emerged as the most discriminative variable between the global distribution of HPs and MPs (Fig. 5), which is consistent with local studies^27–29^ and a literature review^16^. Multiple local studies agree with the prevalence of HP species across different oceanic regions^27,30,31^, as well as with the predominance of MPs in the Arctic^32^. Taken together, the patterns we observed based on metabarcoding data are highly consistent with the literature and support the distribution and diversity conclusions at a larger and global scale.

Despite the global coverage of the *Tara Oceans* data set, a few caveats remain and need to be acknowledged, as each site was sampled only once, and oceanic stations were more numerous than coastal stations. Further, only the most abundant species could be sampled in each sample, with representatives of some major medusozoan taxa missing (*e.g*. Cubozoa) or heavily undersampled (*e.g*. Limnomedusae and Rhizostomae). Additionally, as *Tara Oceans* samples were skewed toward lower sampling depths, we also restricted our analyses to surface and near-surface samples (up to 200 meters). Therefore the conclusions drawn are valid for the euphotic zone, but many medusozoans, notably Coronatae and Siphonophorae^15,33^, inhabit deeper waters, and were not sampled here. It remains to be determined whether the observed patterns will hold with the addition of deep sea data. We are nevertheless confident in the robustness of our results to further sampling of the diversity and distribution of medusozoan. Indeed, not only they largely agree with previous smaller-scale studies, but most importantly they were detected in a variety of oceanic regions and environmental conditions^16,34,35^.

Medusozoans are opportunistic species, feeding on a wide variety of prey, from protists to fish larvae^36^, with both MPs and HPs known to play a major role in the “jelly web”, so called for the ecological and food web role of gelatinous organisms^37^. Medusozoans themselves represent a food source for many species^10^. MPs appeared to occupy a more central position than HPs (Fig. 4A) in a plankton ecological network^23^. Both MPs and HPs types of life cycles co-occurred widely with Alveolata and Rhizaria (Fig. 3B) in the plankton co-occurrence network. Alveolata are known symbionts of many pelagic and benthic cnidarians^38,39^, as well as major food source for many preys of medusozoans. To our knowledge, however, no direct interaction between medusozoans and Rhizaria has been previously reported. This higher centrality implies that MPs tend to associate more strongly with a set of potential biotic interactors. As MPs are found closer to shore, we can hypothesize that the probability of co-occurrence with other organisms is higher than in the open sea and in oligotrophic conditions. The less central position of HPs within the plankton interactome might suggest greater flexibility in terms of biotic interactions and environmental conditions, a likely valuable trait in the rapidly changing and unstable planktonic ecosystem^40^.

In Medusozoa, each independent acquisition of a holoplanktonic life cycle was associated with the colonization of the open ocean, suggesting a direct impact of life cycle evolution on species distribution. We showed that these transitions all occurred during periods of high relative sea level and continental fragmentation. These periods of greater diversity of environments and species isolation globally led to higher rates of animal diversification^41,42^ and thus possibly to higher rates of life cycle evolution. Transitions toward holoplanktonic life cycles in Medusozoa removed the constraints of finding a favorable substrate for settlement, thus providing the opportunity to colonize novel environments. An extension of the duration of the planktonic larval stage was shown in diverse metazoan groups to increase organism dispersal ability allowing them to resist local extinction, decrease inbreeding and competition for resources^2,43^. Marine metazoans also display numerous instances of evolution towards a holoplanktonic life cycle, notably among polychaetes^44^, tunicates^45^, and mollusks^46^. Within these groups, holoplanktonic species often have a cosmopolitan distribution^47,48^ and high ecological impacts^49,50^. Whether similar evolutionary pressures, ecological, and geological contexts favored the loss of the fixed stage and the colonization of the open ocean in the different metazoan groups remains to be investigated.

## Acknowledgments

We thank E. Houliston, F. Lombard, M. Mańko, C. Munro, and C. Sinigaglia, for their feedback on the manuscript and E. Pelletier for help with the omics analyses. This work was funded by the CNRS 80|Prime program (PENELOPE) and supported by the Agence Nationale de la Recherche (ANR-13-PDOC-0016). We wish to thank the CNRS MITI through the interdisciplinary program Modélisation du Vivant (GOBITMAP grant to SC), and the H2020 European Commission project AtlantECO (award number 862923). Tara Oceans (which includes both the Tara Oceans and Tara Oceans Polar Circle expeditions) would not exist without the continuous support of the Tara Ocean Foundation and 23 institutes of research (http://oceans.taraexpeditions.org). We further thank the commitment of the following sponsors: CNRS (in particular the FR2022-GOSEE), European Molecular Biology Laboratory (EMBL), Genoscope/CEA, The French Ministry of Research, and the French Government ‘Investissements d’Avenir’ programmes OCEANOMICS (ANR-11-BTBR-0008). This article is contribution number X of *Tara Oceans*

## Author contributions

L. L and L.G conceived the study, contributed to the analyses and corrected the manuscript. M. B and C.S performed the analyses and wrote the codes and the first version of the manuscript. M.B prepared the figures. O.D.S and S.C helped with the analyses and reviewed the manuscript.

## Methods

### Medusozoan reference phylogenetic tree

All non-redundant ribosomal DNA sequences taxonomically assigned to cnidarians available in the NCBI nucleotide database were retrieved on 22-02-2022. Sequences with a length < 100 nucleotides, unidentified nucleotides rate ≥ 0.01 or described as 16S, 12S, 5S, 60S or whole genome were discarded. A local BLAST database was created from the 27 760 remaining sequences and *Aurelia aurita* complete 18S sequence (AY039208) was aligned by BLAST against this database in order to extract only 18S sequences. Sequences with a blast *p*-value = 0, an alignment length ≥ 1300 bp, and a complete V9 were retained. We made an exception and conserved *Geryonia proboscidalis* 18S, even if the V9 was not complete, in order to have a holoplanktonic representative of the Geryoniidae. Sequences assigned to Myxozoa, or not assigned at the species level, were discarded. Finally, only one sequence per species was selected, selecting the sequence with the longest alignment against the *Aurelia aurita* complete 18S sequence. Very divergent sequences were manually removed (Table S1). 1023 18S ribosomal sequences from cnidarians including 521 sequences from medusozoans were selected. A life cycle type was assigned to each reference sequence using published literature (Table S1). The life cycle of *Poralia rufescens* and the Rhodaliidae *Dendrogramma enigmatica* and *Stephalia dilata* are poorly known and were not defined. All the selected sequences were aligned with mafft L-INS-I^51^ (v7.490). The phylogenomic consensus tree of Kayal et al. (2018)^52^ was used to perform a constrained phylogeny. Only the species present in our alignment were extracted from the consensus tree, and converted to a constraint alignment as described in the FastTree documentation. The 18S phylogenetic tree was then constructed with the topology constrained by the constraint alignment using FastTree with the GTR+CAT model^53^ (v2.1.11). This 18S tree was time-calibrated with calibration points from the literature (Table S2) using the semi-parametric method based on penalized likelihood described by Sanderson (2002)^54^. The implementation available in the ape R package^55^ (chronopl function) was used with a null smoothing parameter, allowing rates to vary as much as possible among branches. The evolution of the life cycle was then simulated 1000 times on the calibrated tree using a stochastic mapping method implemented in the make.simmap function of the R package phytools^56^ (v1.2.0). By taking the posterior density of the stochastic life cycle type mapped at each position of the tree, a total of 6 losses of the benthic stage were found in 81,3% of the simulations. Thus, only the simulations with 6 transitions were selected to date the age of the losses. The average, first and third quartile ages of each loss among the retained simulations were computed.

From the reference tree, we defined 13 medusozoan taxa based on the different transitions from a meroplanktonic life cycle to a holoplanktonic life cycle. Seven meroplanktonic taxa were defined: Anthoathecata (without Porpitidae), Coronatae (without *Periphylla periphylla*), Cubozoa, Leptothecata, Limnomedusae (without Geryoniidae), Rhizostomae, and Semaeostomae (without *Pelagia* spp.). Six holoplanktonic taxa were defined: Geryoniidae, Narcomedusae, *Pelagia noctiluca, Periphylla periphylla*, Porpitidae, Siphonophorae, and Trachymedusae.

### Medusozoan metabarcoding data set processing

The metabarcoding data sets from the *Tara Oceans* expedition (2009-2013, PRJEB6610) and *Tara Polar Circle* (2013, PRJEB9737) were exploited (open access: http://zenodo.org/record/3768510). 781 samples (station/depth/fraction) from 141 stations collected at the surface (SRF) and at the depth of the deep chlorophyll maximum (DCM) were explored. The following size fractions were included in the analysis: “0.8 to 5 μm and 0.8 to 2000 μm,” “3 to 20 μm and 5 to 20 μm,” “20 to 180 μm,” and “180 to 2000 μm”. In order to have a homogeneous data set, we considered the fraction 3-20 μm, mainly used during the TARA Polar Circle expedition, equivalent to the fraction 5-20 μm used during the*Tara Oceans* expedition, as well as fractions 0.8-5 μm equivalent to 0.8-2000 μm, as samples showed similar diversity and community composition^23^. In the case where both size fractions were available for the same sample, the more ubiquitous 0.8-5 μm size fraction was selected. For the sole station TARA_124_SRF for which the two sizes 3-20 μm and 5-20 μm were sampled, only the latter one was retained.

For each sample, the V9-18S ribosomal DNA from Ibarbalz et al. (2019)^21^ was analyzed. Data processing has been previously described in de Vargas et al. (2015)^57^ and consisted of a quality check, a filtering of the metabarcodes present in at least 2 samples with more than 3 reads followed by clustering into operational taxonomic units (OTUs) using the Swarm software^58^. Each OTU was then taxonomically and functionally annotated by an optimal global aligner (VSEARCH v2.3.1^59^) against an updated version of the PR2_V9, an SSU V9 rDNA reference database available at http://zenodo.org/record/3768951. To determine the type of life cycle of each medusozoan OTU (*e.g*. meroplanktonic or holoplanktonic), an assignment at the genus, family, or order level was needed. To achieve this, the 2 796 OTUs assigned to cnidarian were extracted from the *Tara Oceans* data set. The representative sequence of each OTU was aligned by BLASTn against the PR2_V9 database enriched in a non-redundant manner with Opisthokonta 18S sequences with a length between 100 and 10 000 pb extracted from the nucleotide database of the NCBI. This step allowed the elimination of OTUs falsely assigned to Cnidaria. OTUs whose first hit was assigned to Cubozoa, Scyphozoa, or Hydrozoa classes with an identity percentage above 80% were retained, which represented 1 933 OTUs. We removed OTUs assigned to Staurozoa (1 OTU) because they exhibit a mainly benthic life cycle^60^ and Myxozoa (13 OTU) because of their parasitic life cycle^61^.

A taxonomic identity was assigned to the selected sequences by a phylogenetic approach: V9 sequences of the 1 933 medusozoans OTUs were placed in the 18S constrained reference tree (see above) using pplacer^62^ (v1.1.alpha19). For each OTU, all placements with a wanted rank at the species level were retained and the common ancestor node of these placements was selected to assign a taxon to the OTUs. In cases where the placement by pplacer was not accurate enough to infer a defined taxon (see above) or a life cycle, the blast assignment of OTUs was retained if the percentage of identity was higher than 97%. According to the taxa assigned, the type of life cycle was determined for each OTUs. In the case of Coronatae, the 18S V9 rRNA of *Periphylla periphylla* (holoplanktonic) and *Paraphyllina ransoni* (meroplanktonic) is not discriminative enough to distinguish these two species, and those OTUs were removed. Finally, OTUs assigned to Staurozoa or Anthozoa and those whose assignment did not allow to determine a life cycle or a taxon were removed. Taxonomic assignments by blast and pplacer were compared and OTUs with conflicting taxon assignments were eliminated. These steps excluded 151 OTUs leading to 1782 OTUs properly assigned (Fig. S1A). In order to remove possible false positive OTUs, only OTUs with a maximum relative abundance > 0.0015% and found in at least 2 samples (station and depth, all size fractions) were selected. The threshold was defined visually by making a histogram of the maximum relative abundance of OTUs, and selecting a threshold that removes a large number of very low abundant OTUs. It eliminated 0.06% of the reads while eliminating 80.6% of the OTUs. No Cubozoa were retained after the filtration. 300 OTUs were conserved.

### Environmental data selection

The publicly available environmental data sets containing the water column features at the sampling location (https://doi.pangaea.de/10.1594/PANGAEA.875579) and the mesoscale features (https://doi.pangaea.de/10.1594/PANGAEA.858201) collected during the *Tara Oceans* (2009-2013) and *Tara Polar Circle* expedition (2013) were used in this study. The strategy and methodology employed during the campaign are described in Pesant et al. (2015)^18^. A meaningful environmental data set was extracted from the variable available and homogenized by replacing the missing values with the median value of the oceanic region defined by Longhurst (2010)^63^ in order to perform redundancy analysis (RDA) which cannot handle missing values. The table contained environmental variables for 781 of our 785 samples. The 4 samples of the station number 011, lacking environmental information, were not included in the statistical analyses (Table S5).

### Statistical analysis

Data analysis and statistical tests were performed using R (v4.2.1)^64^ with the packages vegan 2.6.2^65^, tidyverse 1.3.2^66^, and custom scripts. Rarefaction curves were performed using 100 Monte Carlo resampling using the package MonteCarlo 1.0.6^67^. Analysis of the rarefaction curves revealed that HP OTUs are more diverse than MP OTUs when the analysis is performed on a small number of samples (Fig. S3). This may be due to the fact that there are more different MP OTUs than HP OTUs but the sampling and sequencing depth required to detect all MP OTUs present in the environment is higher than the one required to detect all HP OTUs. To avoid underestimating MP diversity, we average the relative abundance of OTUs in the samples from the same station (depth/fraction). All the other analyses were carried out by samples (station/depth/fraction). For the RDA, the environmental data set was used as an explanatory matrix and the OTU count matrix as a response matrix. The relative abundances of OTUs from each medusozoan were summed for each sample and normalized using a Hellinger transformation^68^. Environmental data were centered prior to the analysis and then selected by stepwise model selection using permutation with the ordistep function from the vegan package (options: 1000 steps). The model selected 10 environmental response variables (bathymetric depth, acCDOM, density, nitrate, salinity, PAR, residence time, depth nominal, bbp470 and oxygen).

### Co-occurrence network analysis

The global ocean plankton community interactome published by Chaffron et al. (2021)^23^ was used to investigate the co-occurrence between meduzosoan OTUs and other OTUs. Only edges with positive correlations were selected. We used our taxonomic assignments for medusozoan nodes and classified the assignments of non-medusozoan nodes into larger taxa: Alveolata, Anthozoa, Bacteria, Copepoda, Metazoa, Rhizaria, Stramenopiles, Tunicata, and others. Interactome analysis was performed using the packages igraph (1.3.4)^69^ and ggraph (2.0.6)^70^.

### Phylogenetic regression

An OTU phylogenetic tree was created by adding the OTUs in the calibrated 18S reference tree (see the “Medusozoan reference phylogenetic tree” section) at the node corresponding to their taxonomic assignment, and then removing the reference sequences to only conserve the OTUs. The evolution of the colonized bathymetric depth and the acCDOM was estimated by mapping their median values onto the OTUs phylogenetic tree using the anc.ML function from the phytools^56^ (v. 1.2.0) R package. Finally, the correlations between the presence of a benthic stage in the OTU life cycles and the median bathymetric depth or acCDOM were tested by a phylogenetic logistic regression as described in Ives and Garland (2010)^71^ using the R package phylolm (V. 2.6.2)^72^.

---

**Supplementary Fig. 1.**
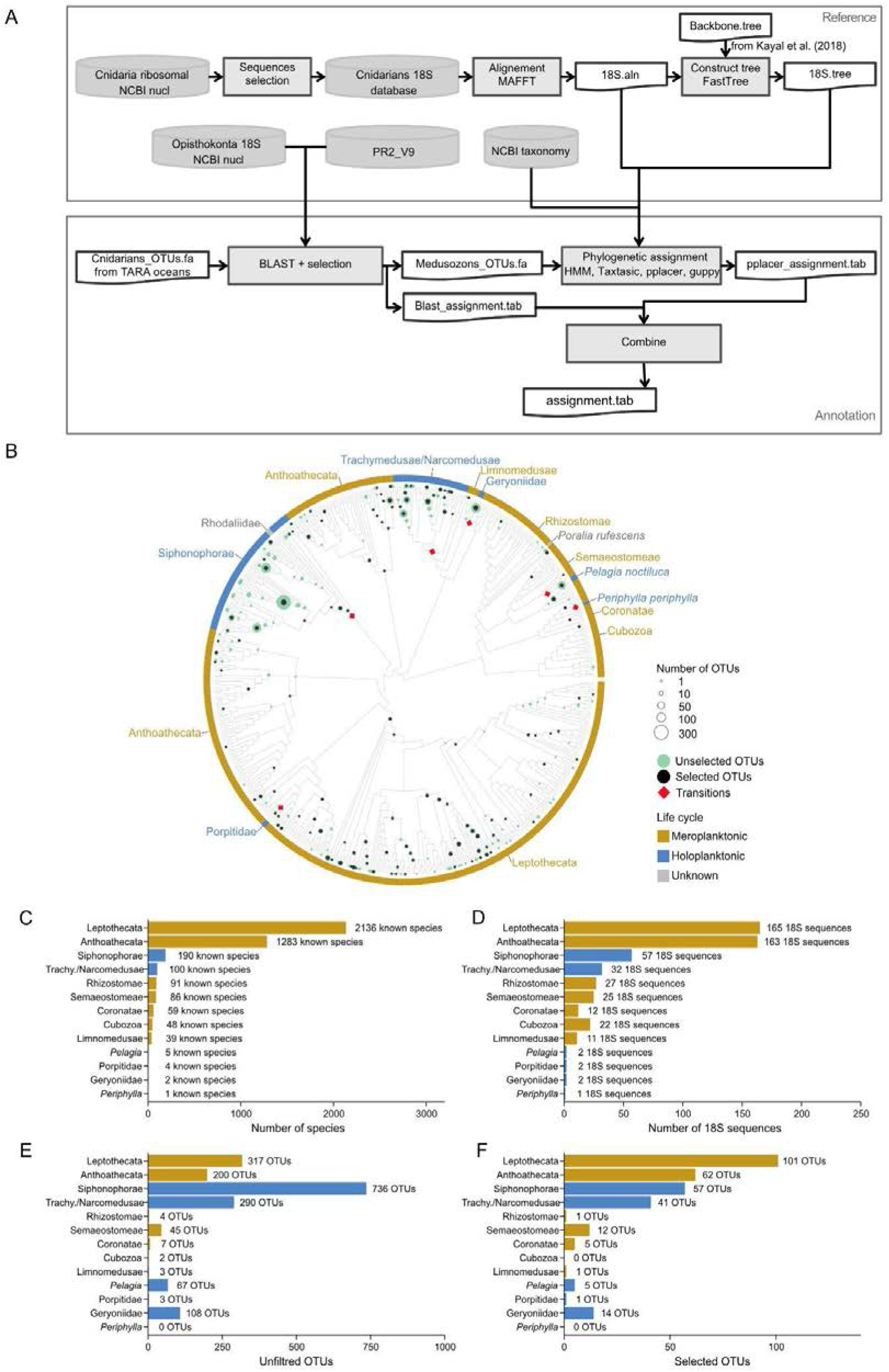
Taxonomic assignment of medusozoan OTUs. (A) Schematic representation of the pipeline used to assign medusozoan OTUs. (B) Phylogeny inferred on the 18S sequence using FastTree with a constraint tree from Kayal et al. (2018)^52^. The gray branches correspond to all 18S reference sequences included in the phylogeny. OTUs have been placed in the phylogeny using pplacer. Selected OTUs (maximum relative abundance > 0.0015% and found in at least 2 samples) are represented by black dots; unselected OTUs are represented by green dots. The size represents the number of OTU found at each branching point. The outer circle refers to the type of life cycle for each species, yellow for meroplanktonic, blue for holoplanktonic and gray when it is unknown. Transitions from a meroplanktonic to a holoplanktonic life cycle are represented by the red squares (C) Number of known species for each medusozoan taxa extracted from the World Register of Marine Species (WoRMS)^22^. (D) Number of 18S sequences for each medusozoan taxa used for the reference medusozoan phylogeny. (E) Number of OTUs assigned to each medusozoan taxa before filtering on their relative abundance and the number of stations they are found. (F) Number of selected OTUs for each medusozoan taxa.

**Supplementary Fig. 2.**
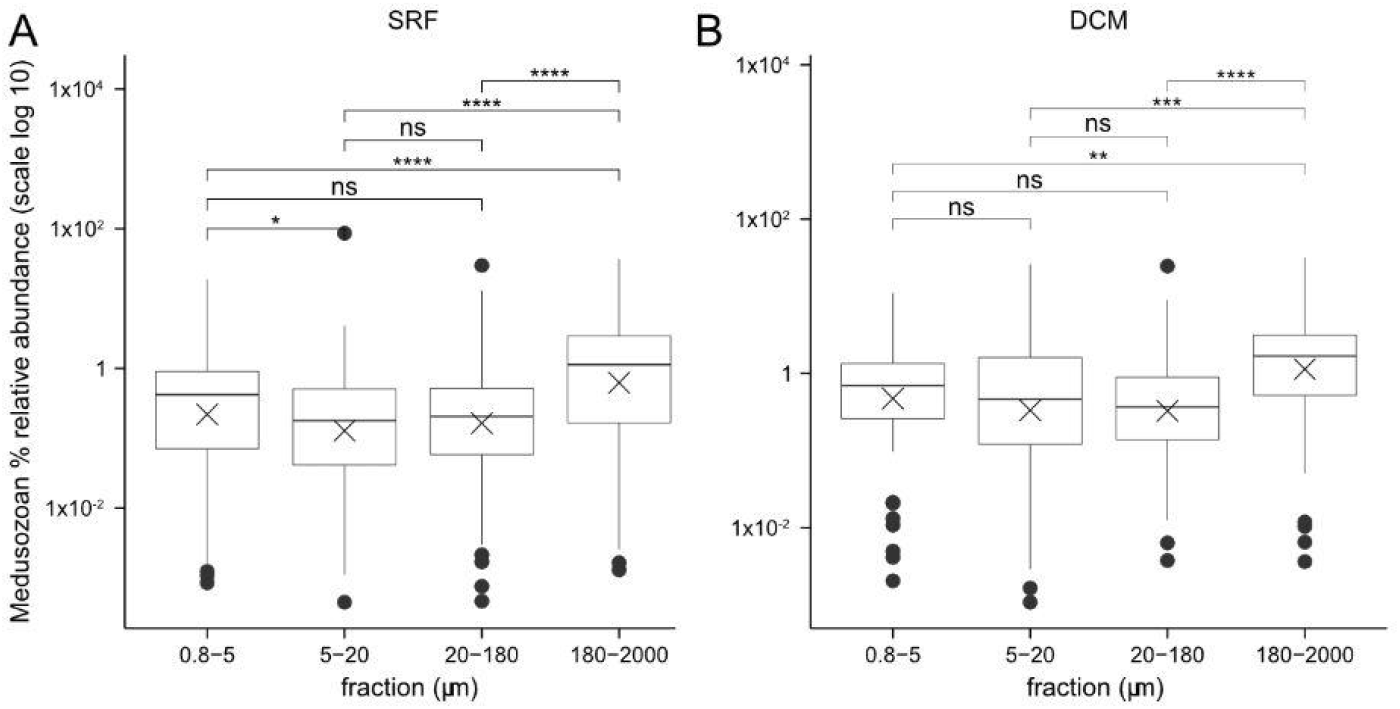
Relative abundance of medusozoan OTUs as a function of the size fraction for the surface (SRF, A) and the deep chlorophyll maximum layer (DCM, B) samples. *P*-values were computed using Wilcoxon test, **** *p*-value <= 0.0001; *** *p*-value <= 0.001; ** *p*-value <= 0.01; * *p*-value <= 0.05; ns not significant.

**Supplementary Fig. 3.**
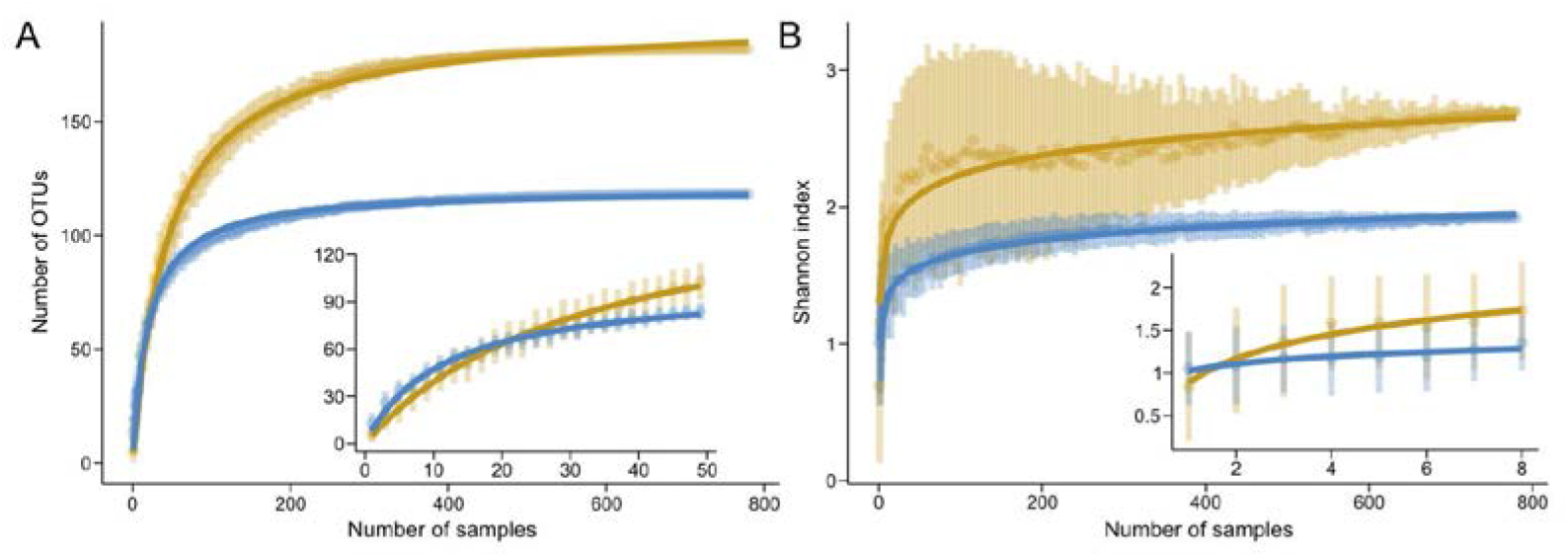
Rarefaction curve for holoplanktonic and meroplanktonic medusozoans based on the number of OTUs (A) and the Shannon index (B). Insets show the rarefaction curve performed on fewer samples for both the number of OTUs and the Shannon index. Both metrics showed saturation at the global scale, unless the rarefaction analysis was done on a small number of samples (e.g eight samples). In this latter case, HP OTUs represent a greater richness and diversity.

**Supplementary Fig. 4.**
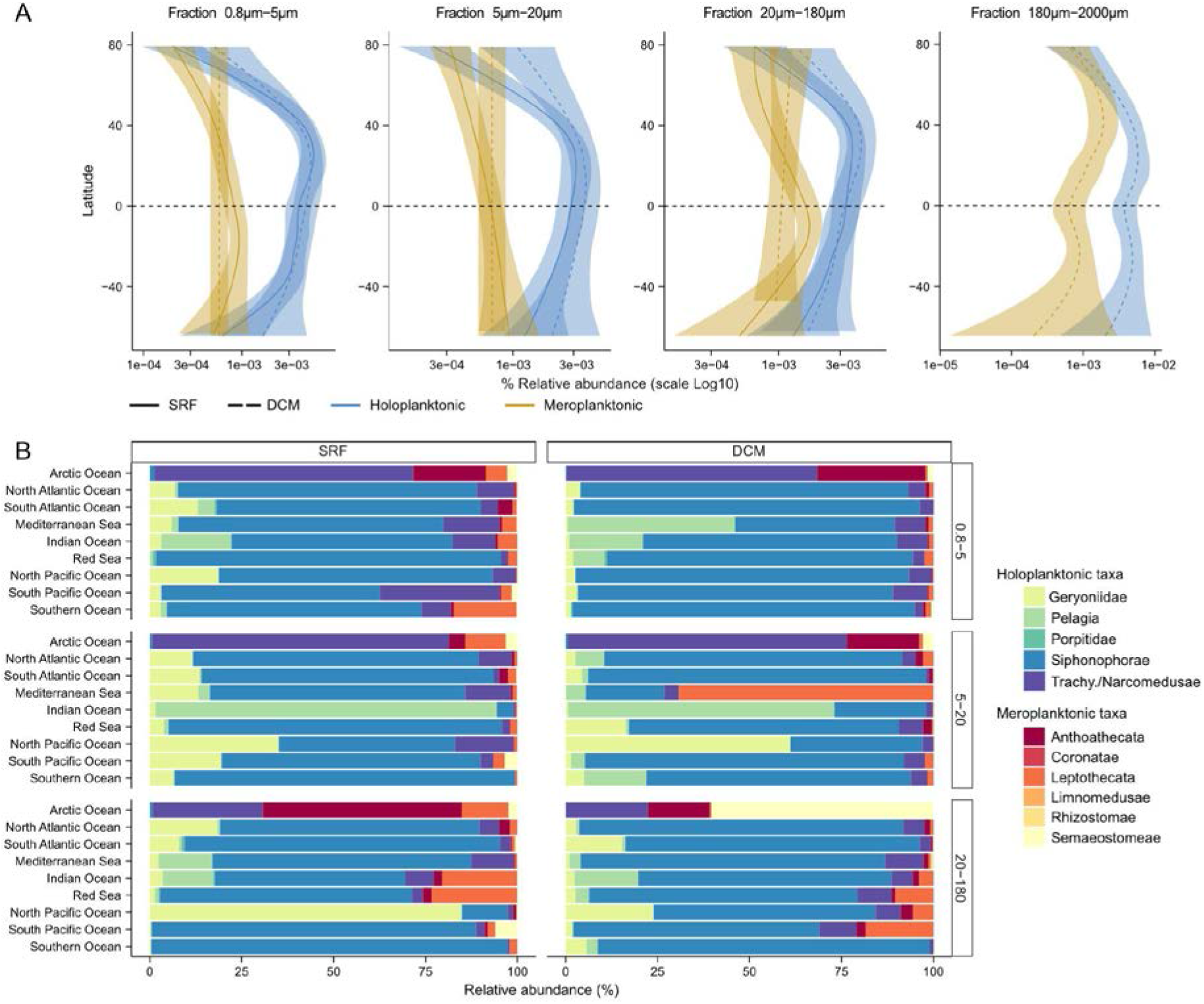
Distribution and diversity of medusozoans for the size fractions not shown in Fig. 3. (A) Latitudinal gradient of relative abundance for both types of life cycle with the x-axis log10 scale. Solid and dotted lines correspond respectively to SRF and DCM samples. (B) The proportion of relative abundance of OTUs assigned to the different medusozoan taxa according to Longhurst biomes (Longhurst, 2010)^63^ for the size fractions not shown in Fig. 3.

**Supplementary Fig. 5.**
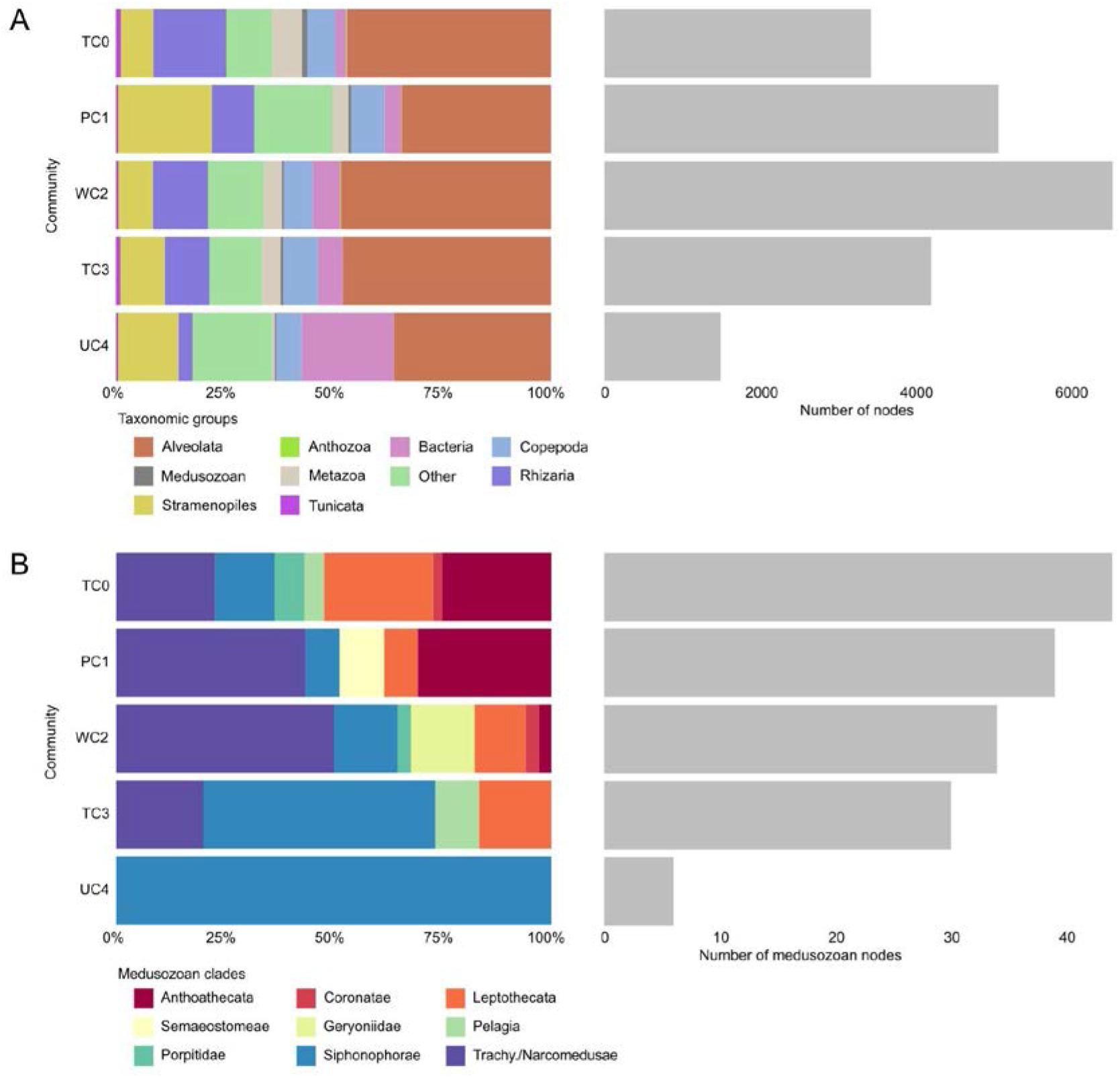
Non medusozoan and medusozoan taxa composition in the co-occurrence network communities. (A) Taxonomic composition and the number of nodes found in each community (B) Medusozoan taxa composition and the number of nodes found in each community.

**Supplementary Fig. 6.**
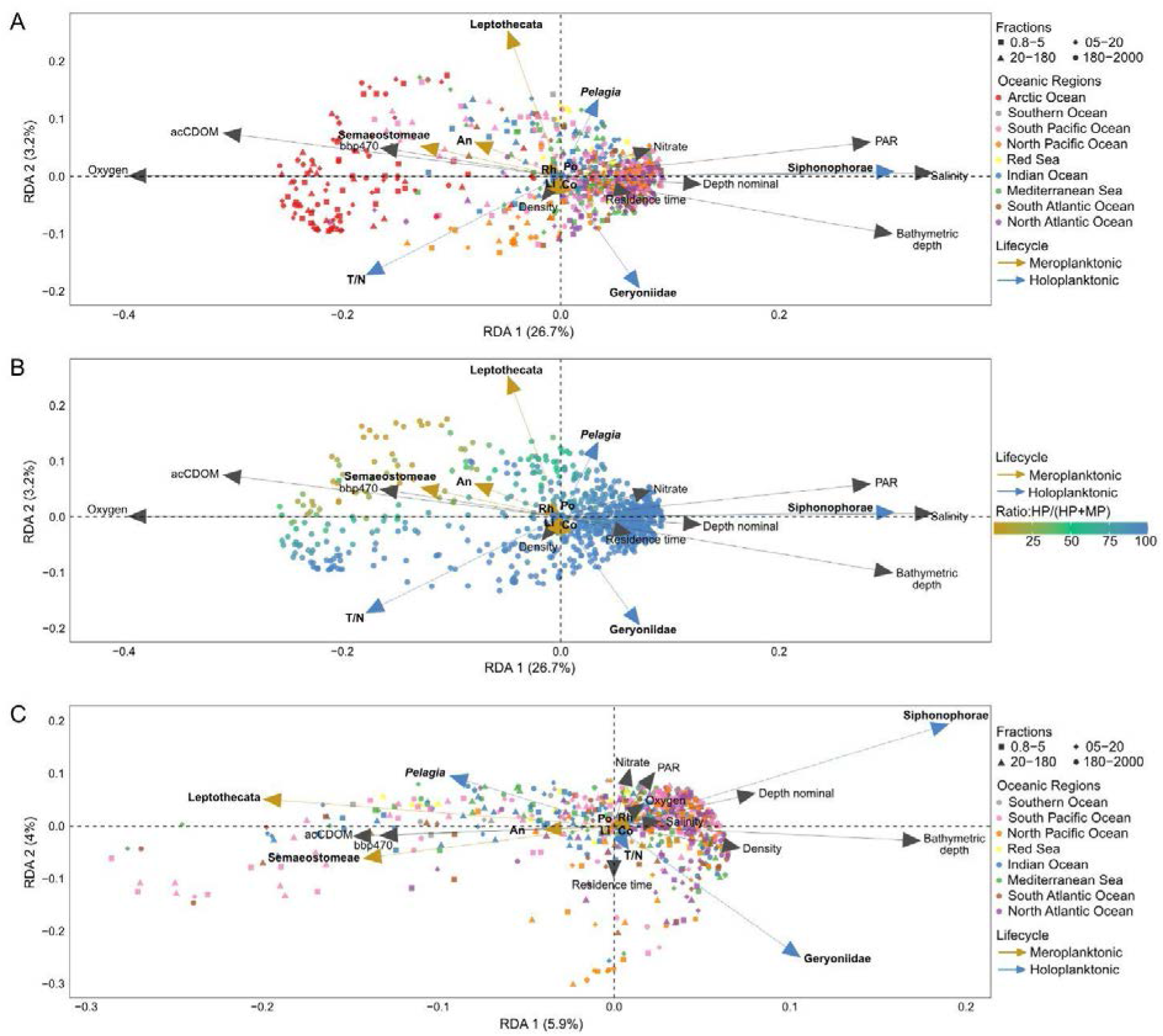
Impact of environmental variables on medusozoan distribution with and without Arctic samples. (A) Triplot of the redundancy analysis (RDA) with Arctic samples, dots are colored by their oceanic region. (B) Triplot of the redundancy analysis (RDA) with Arctic samples, dots are colored by the life cycle ratio (abundance of holoplanktonic OTUs on total medusozoan abundance) so that the blue dots represent samples dominated by holoplanktonic OTUs and the yellow dots samples dominated by meroplanktonic OTUs. (C) Triplot of the redundancy analysis (RDA) without Arctic samples showing the explanatory environmental variables in black, and the response variable, the relative abundance of medusozoan clades, in blue for holoplanktonic taxa and yellow for meroplanktonic taxa. Each dot corresponds to a sample (i.e. one station at one depth and one size fraction) and is colored by its oceanic region. Abbreviations: An Anthoathecata, Co Coronatae, Li Limnomedusae, Po Porpitidae, Rh Rhizostomae, T/N Trachymedusae / Narcomedusae.

**Supplementary Fig. 7.**
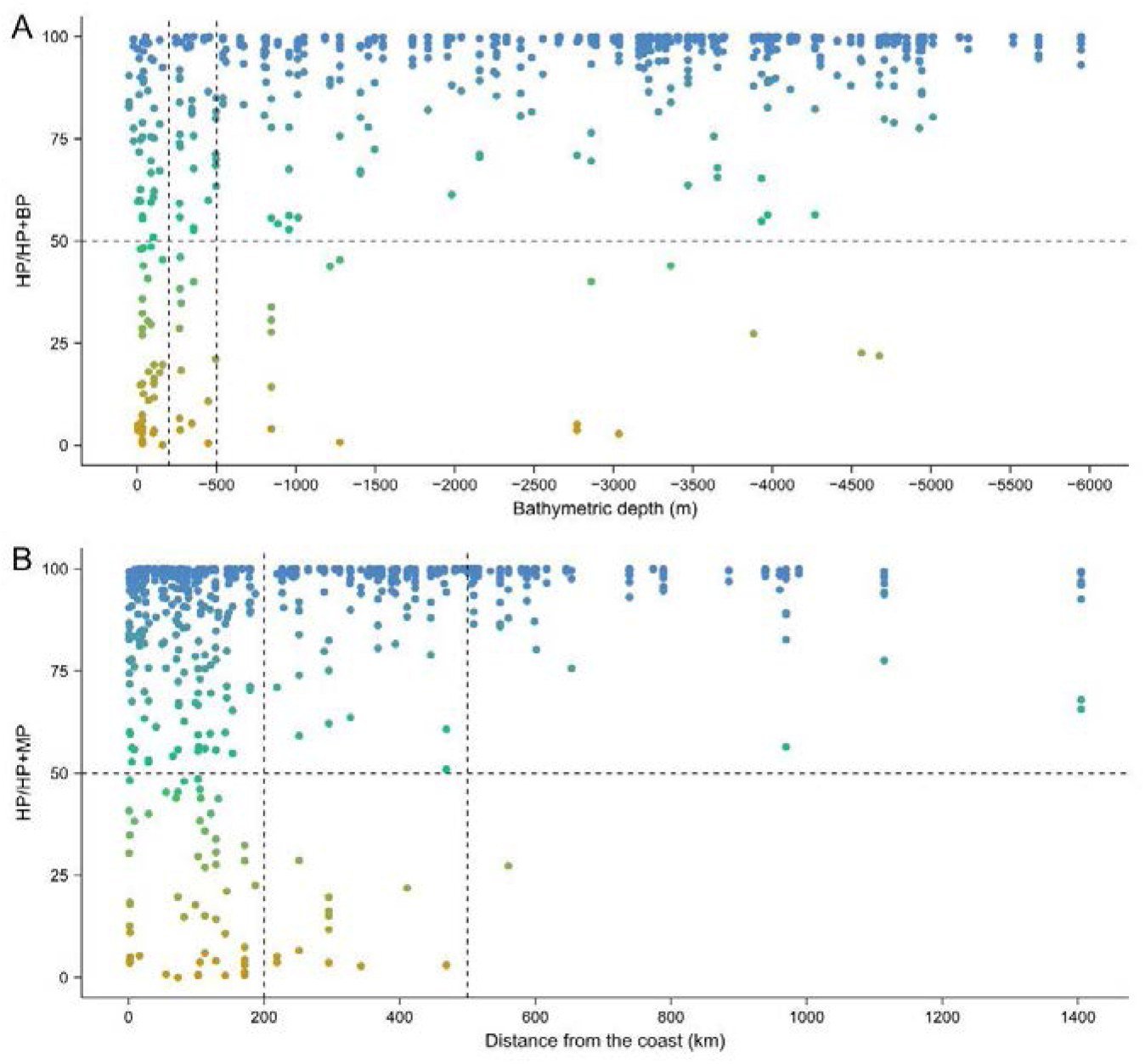
Relationship between the life cycle ratio in the samples (including Arctic samples) and the bathymetric depth (A), or the distance from the coast (B). The vertical lines correspond to bathymetric depth of 200 meters and 500 meters and distances from the coast of 200 kilometers and 500 kilometers respectively.

**Supplementary Fig. 8.**
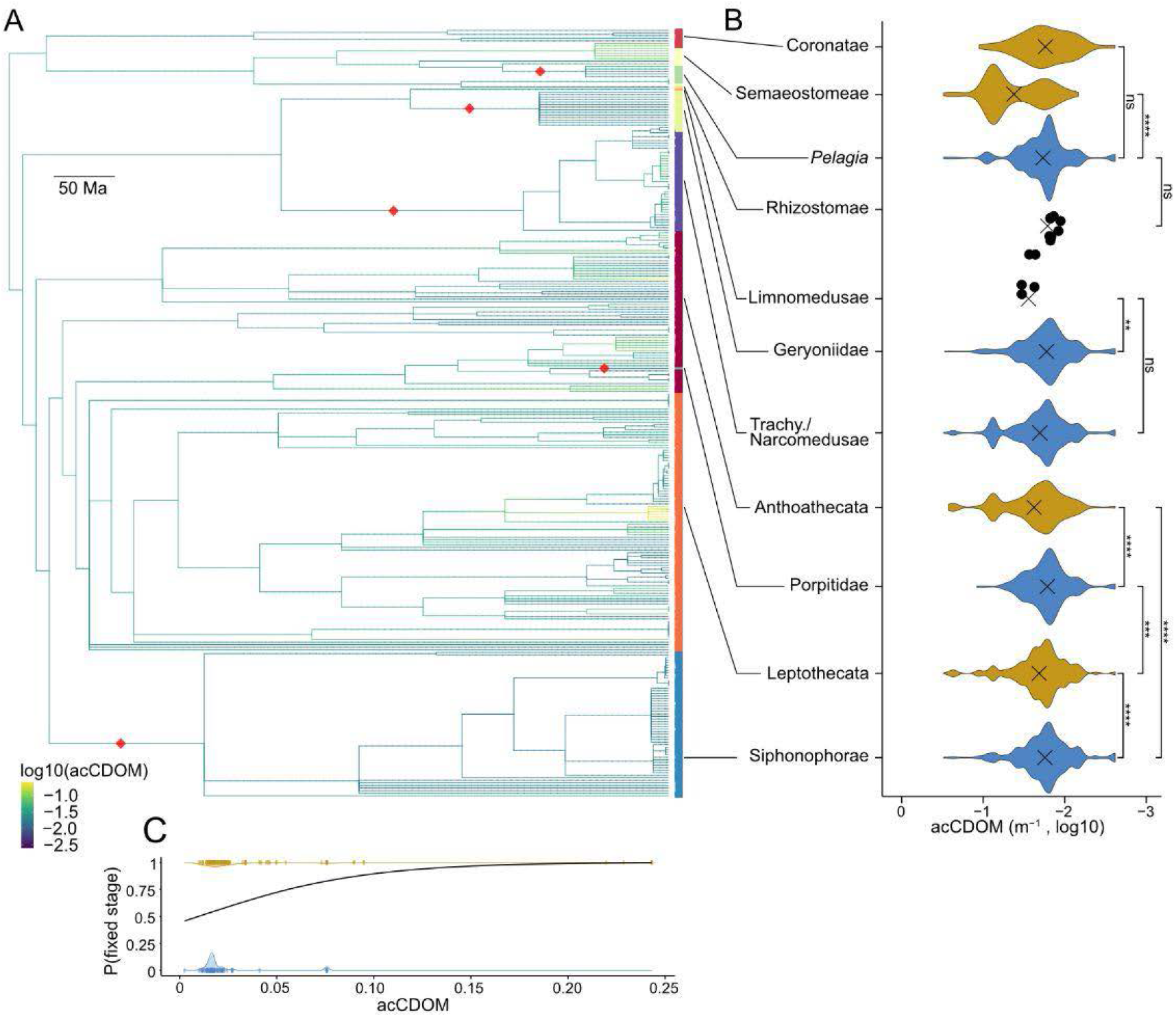
Phylogenetic relationship between medusozoan life cycles and acCDOM (turbidity; in meters^-1^). (A) Phylogenetic tree of OTUs on which the evolution of the median value of the acCDOM (log10) has been estimated using maximum likelihood and used to color the branches of the tree. The red square illustrates transitions from a meroplanktonic life cycle to a holoplanktonic life cycle. (B) Violin plot reporting the acCDOM values by OTUs per taxa. *P*-values were computed using Wilcoxon test, **** *p*-value <= 0.0001; *** *p-* value <= 0.001; ** *p*-value <= 0.01; * *p*-value <= 0.05; ns not significant.. (C) Phylogenetic logistic regression showing the relationship between acCDOM and the type of life cycle. *P*-value: 2.5 × 10^-5^.

## Supplementary tables

Supplementary Table 1: 18S references included in the reference tree and their life cycle according to the literature. [separate file]

**Supplementary Table 2:** Calibration points.

| Clade | Earliest fossils (Ma) | Reference |
| --- | --- | --- |
| Cnidaria | 560 | Dunn et al. (2022)<sup>1</sup> |
| Anthozoa | 540 | Han et al. (2010)<sup>2</sup> |
| Medusozoa | 535 | Wang et al. (2022)<sup>3</sup> |
| Scyphozoa | 505 | Cartwright et al. (2007)<sup>4</sup> |
| Leptothecata | 470 | Cope (2005)<sup>5</sup> |
| Macrocolonia | 380 | Song et al. (2021)<sup>6</sup> |
| Stylasteriidae | 65 | Lindner et al. (2008)<sup>7</sup> |
| Oculina | 56 | Quattrini et al. (2020)<sup>8</sup> |

Supplementary Table 3: OTU assignment and their life cycle. Available upon request. [separate file]

**Supplementary Table 4:** Summary of the distribution of medusozoan OTUs and reads among oceanic region, depth and size fraction.

| Ocean region | Number of OTUs | Number of reads |
| --- | --- | --- |
| Arctic Ocean | 80 | 1 664 018 |
| North Atlantic Ocean | 144 | 1 692 887 |
| South Atlantic Ocean | 128 | 795 981 |
| Mediterranean Sea | 146 | 3 152 364 |
| Indian Ocean | 185 | 5 474 878 |
| Red Sea | 106 | 464 021 |
| North Pacific Ocean | 130 | 2 987 277 |
| South Pacific Ocean | 198 | 2 341 954 |
| Southern Ocean | 49 | 262 958 |
| Depth | Number of OTUs | Number of reads |
| SRF | 295 | 11 328 654 |
| DCM | 253 | 7 507 684 |
| Fraction (µm) | Number of OTUs | Number of reads |
| 0.8-5 | 268 | 2 448 747 |
| 05-20 | 236 | 4 594 320 |
| 20-180 | 284 | 3 172 645 |
| 180-2000 | 267 | 8 620 626 |

Supplementary Table 5: Holoplanktonic OTUs ratio (HP/(HP+MP relative abundance)) across samples. [separate file]

